# A light-weight, data-driven segmentation method for multi-state Brownian trajectories

**DOI:** 10.1101/2025.06.05.658053

**Authors:** Ismail El Korde, Jason M. Lewis, Erik Clarkson, Tommy Dam, Peter Jönsson, Tobias Ambjörnsson, Joakim Stenhammar

## Abstract

Single-particle tracking methods have emerged as a crucial tool for the characterization of dynamical and diffusive processes in a range of biological and synthetic systems. Here, we propose a simple and light-weight yet accurate method for the segmentation of multi-state Brownian trajectories based on an optimised Gaussian filtering of the displacement time series combined with an automated fitting to a Gaussian mixture model. We verify our method using synthetic, 2-state Brownian trajectories and show that our method provides high levels of accuracy in terms of segmentation and the estimation of self-diffusion coefficients for reasonably well-separated values of the diffusion coefficients. We furthermore demonstrate the feasibility of our method on experimental systems using single-particle tracking data for diffusing membrane proteins bound to a supported lipid bilayer. Compared to methods based on deep learning or hidden Markov models our method imposes a much lower computational load, making it suitable for fast and accurate online processing of single-particle trajectories from microscopy images.

## Introduction

The revolution in optical microscopy techniques over the last three decades have significantly enhanced the ability to investigate biological systems at the molecular level. Among these, single-molecule fluorescence microscopy [1–3] in conjunction with single-particle tracking (SPT) algorithms [4–10] has revolutionised the study of molecular dynamics in living cells by providing unprecedented spatiotemporal resolution. By enabling the tracking of individual molecules, these methods have yielded crucial insights into intracellular transport, protein-protein interactions, and the fundamental mechanisms that govern cellular processes [1, 7, 11–16], as well as fundamental knowledge about diffusive processes [17, 18].

A fundamental challenge in SPT analysis is the characterisation of multiple diffusive states based on single-particle trajectories, which can often exhibit significant heterogeneity due to complex interactions with the cellular environment [16, 19–22]. The primary biophysical example of such heterogeneity is the change in diffusion constant of a ligand molecule when bound to a substrate, with two recent examples being the binding of the Trigger Factor chaperone protein binding to the ribosome surface [16] and the transient binding of membrane-associated immune cell proteins to their corresponding receptor proteins [25]. The binding between a molecule and a receptor typically leads to a radical reduction in the diffusion coefficient of the complex compared to that the freely diffusing ligand, and the ability to quantitatively analyse such multi-state diffusive processes in terms of their switching times and diffusion coefficients is thus essential for extracting parameters such as protein-ligand binding affinities, which eventually control important biological processes such as cellular signaling [23–25]. The traditional approach for analysing diffusive processes in homogeneous systems involves the calculation of the mean-square displacement (MSD), which, for molecules undergoing simple Brownian motion characterised by a single diffusion coefficient, serves as a straightforward method for inferring molecular transport properties [26–28]. For biological systems with multi-state or spatially heterogeneous diffusion, MSD-based analysis methods are however less straightforward, since a temporally and spatially resolved sampling of the MSD is very statistically challenging [20, 29].

To address this limitation, more sophisticated methods have been developed to achieve parameter estimation and segmentation of multi-state diffusive trajectories. One widely used class of methods is based on so-called *hidden Markov models* (HMMs), whereby trajectories are segmented based on probabilistic modelling to find the maximum-likelihood (or Bayesian) estimate of the diffusion coefficients *D*_*i*_ and transition probabilities *p*_*i j*_ of the different states [30–39]. Traditionally, HMM models require as input a predefined number of diffusive states [32], although more recent HMM-based methods overcome this requirement, instead inferring the most probable number of diffusive states directly from the data [30, 33–35, 38]. Although significantly more accurate than MSD-based methods, inferring diffusive parameters using HMM methods can become computationally demanding for complex systems with many diffusive states. In addition to these statistical methods, several well-performing deterministic methods for change-point detection between diffusive states have been developed [40–44], together forming a large toolbox of segmentation algorithms for multi-state particle trajectories.

More recently, methods based on machine learning (ML) [45, 46] and, more specifically, deep learning [47–53] have emerged as a versatile and accurate tool to handle the analysis of single-molecule dynamics, including extraction of trajectories from (potentially noisy) microscopy images, segmentation of multi-state Brownian trajectories, and the analysis of “anomalous”, non-Brownian diffusion processes, where the MSD grows super-or sublinearly with time [47, 48, 50, 51]. While these data-driven approaches often demonstrate impressive accuracy in analysing complex diffusion behaviors, they are typically computationally heavy and require large training datasets. Furthermore, they typically lack physical interpretability due to the complex nature of the ML architecture, making it difficult for the user to assess the quality of the parameter extraction.

In this study, we propose an alternative, light-weight approach for the segmentation of multi-state Brownian trajectories. Our method uses a simple, one-dimensional Gaussian filter that acts on the displacement time series to achieve efficient noise reduction. The resulting displacement distribution is fitted to a Gaussian mixture model in order to adjust the filter width in an automated manner to optimally classify the different diffusive states. In spite of its relative simplicity and low computational cost, we show that, for reasonably well-separated and long-lived diffusive states, the methodology is able to achieve accurate segmentation while not relying on the large amounts of (synthetic or experimental) training data required for ML-based methods. Compared to these methods, our approach is furthermore physically transparent, in that the automated fitting and filter optimisation process can be monitored to evaluate the quality of the segmentation. In contrast to HMM-based approaches, segmentation of the trajectory is done *before* parameter estimation, meaning that diffusion constants and lifetimes can be estimated using the well-established methods for single-state trajectories, as we will demonstrate below. In summary, our method adds to the growing toolbox of multi-state SPT algorithms, providing a simple and light-weight tool for the segmentation of single-particle trajectories and the estimation of self-diffusion coefficients and state lifetimes.

## Results

Throughout this paper, we will consider the trajectory of a single Brownian particle which can adopt a number of discrete diffusive states *i*, each characterised by a self-diffusion coefficient *D*_*i*_. The different states can, for example, model the binding and unbinding of a ligand to a receptor or the clustering between two protein molecules in a lipid bilayer, leading to a sudden decrease in diffusivity. The transition between states is described as a Poisson process with transition rates 𝜆_*i j*_, representing the probability per unit time of transitioning from state *i* to state *j*. For simplicity of presentation, our analysis will be restricted to the case of 2-state Brownian diffusion in 2 spatial dimensions, but the method can be generalised to higher-dimensional trajectories and/or higher numbers of states in a straightforward manner.

### Model description

Synthetic, 2-state Brownian trajectories are generated by solving the discretised, overdamped Langevin equation using Euler’s method:

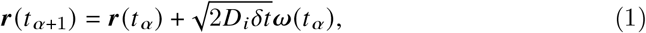

where *t* _𝛼_ = 𝛼𝛿*t* is the discretised time, 𝛿*t* is the time step used to generate the trajectory (not to be confused with the microscope acquisition time, Δ*t*, which we introduce below), and 𝝎 is a vector with the dimension of the system where each component is drawn from a normal distribution with zero mean and unit variance.

At each time step the particle changes its diffusion coefficient from *D*_1_ (resp. *D*_2_) to *D*_2_ (resp. *D*_1_) with a predefined transition rate 𝜆_12_ (resp. 𝜆_21_) with the transition probability [31]

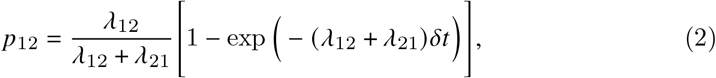

where *p*_11_ = 1 − *p*_12_ is the probability of remaining in state 1, and *p*_21_ is obtained by interchanging the indices 1 and 2 in Eq. (2). The average lifetimes are thus given by 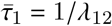 and 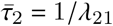.

To mimic the blur induced by the camera’s shutter count, following Ref. [54], we introduce the acquisition time Δ*t* ≡ *n*𝛿*t*, where the integer *n* is the number of discrete time steps over which the image is acquired. We assume constant illumination throughout the acquisition time Δ*t*, meaning that all positions contribute equally to the average position measured over the time window. To mimic this process, we first generate a Brownian trajectory of length *T* by solving Eq. (1), which we then divide into *T*/*n* non-overlapping segments of length *n*, separated by Δ*t*. The blurred trajectory is constructed by taking the arithmetic mean of the particle’s position within each segment. In addition, we mimic the errors introduced by limited spatial resolution and the subsequent tracking process by adding a Brownian noise 𝝃 of zero mean and variance 𝜎^2^ [37]. Taken together, the noisy trajectory 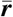 is thus obtained as

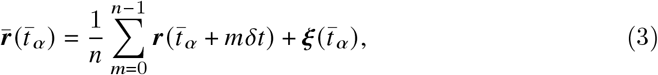

where 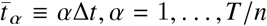 now represents time measured in units of the camera acquisition time. This procedure introduces both temporal and spatial correlations into the trajectory, which must be considered when analysing the particle’s dynamics, such as determining the diffusion constant [54] or transition probabilities. An example of such a trajectory is presented in Fig. 1, illustrating the motion before and after the incorporation of motion blur and localization error. In the following, we will use Δ*t* as our basic time unit when presenting our data, unless stated otherwise.

**Fig. 1.**
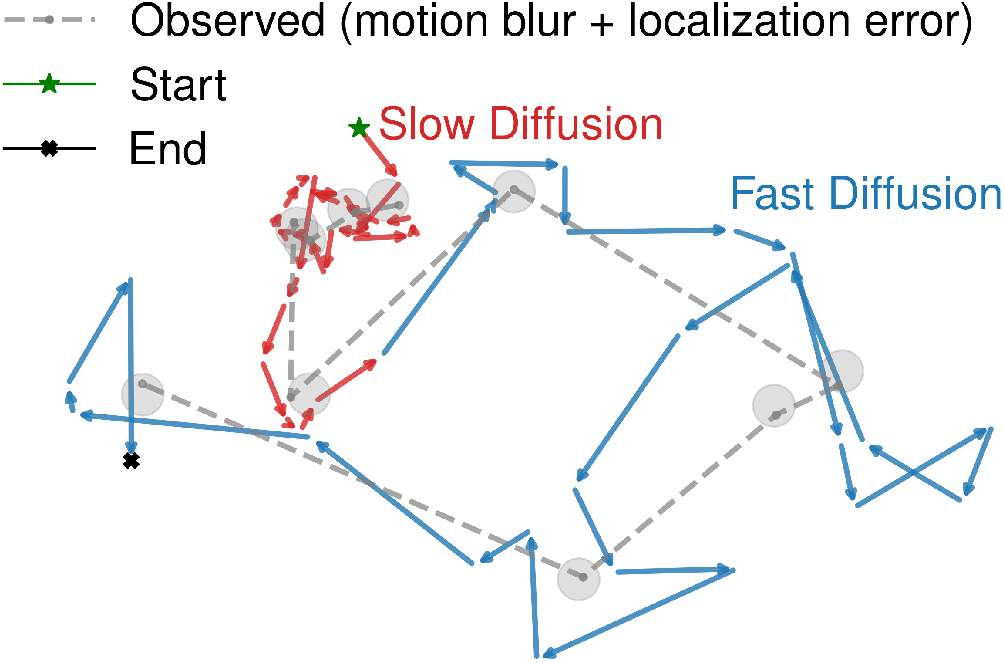
Example of a 2-state Brownian trajectory: The true trajectory is represented by color-coded arrows, while the observed trajectory, shown as a dashed gray line with circular markers, is affected by motion blur (*n* = 5), where the path is averaged over non-overlapping time intervals. Localisation error is simulated by adding Gaussian noise to the blurred positions, represented by circles around each point, reflecting the uncertainty in position measurement.

### Segmentation algorithm

The primary goal of the method is to infer the state of the particle at each time step based on its positional time series and the expected number of states. As schematically depicted in Fig. 2, we start by computing the displacement time series from the trajectory 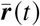, as generated from Eq. (3), by computing the Euclidean norm of the distance Δ*r* between two successive positions of the particle:

**Fig. 2.**
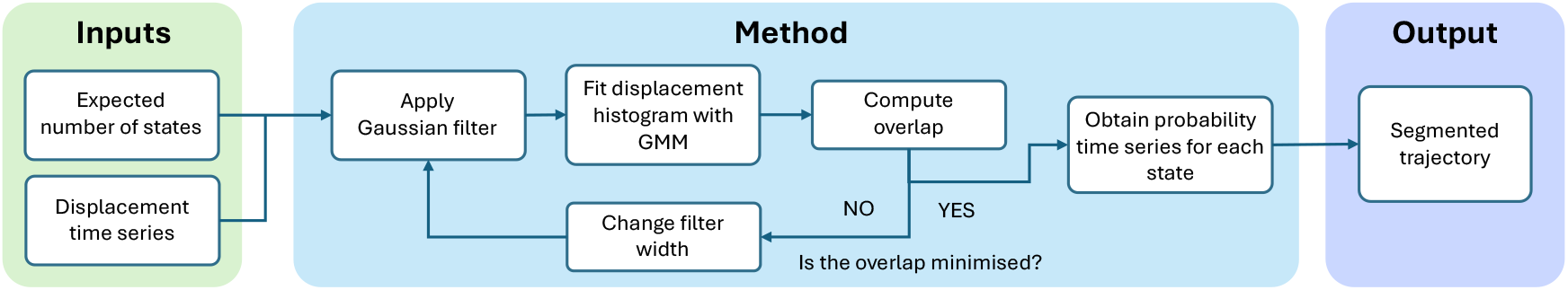
Schematic of the different steps in the method: Given a set of trajectories and an expected number of states, the overlap between the displacement distributions is minimised using an optimised Gaussian filter acting on the displacement time series. The output consists of a segmentation of the given trajectory, inferring the current state of the particle at each given time point.

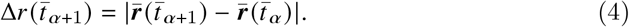

Fig. 3a shows an example of such a noisy time series, clearly showing a large spread across the two states. Since the displacements Δ*x* and Δ*y* in each dimension are normally distributed with mean 𝜇_*i*_ = 0 and variance 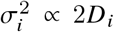, the displacement probability distribution *P*_*i*_ (Δ*r*) in state *i* of 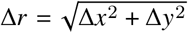 is Rayleigh distributed, *i*.*e*.

**Fig. 3.**
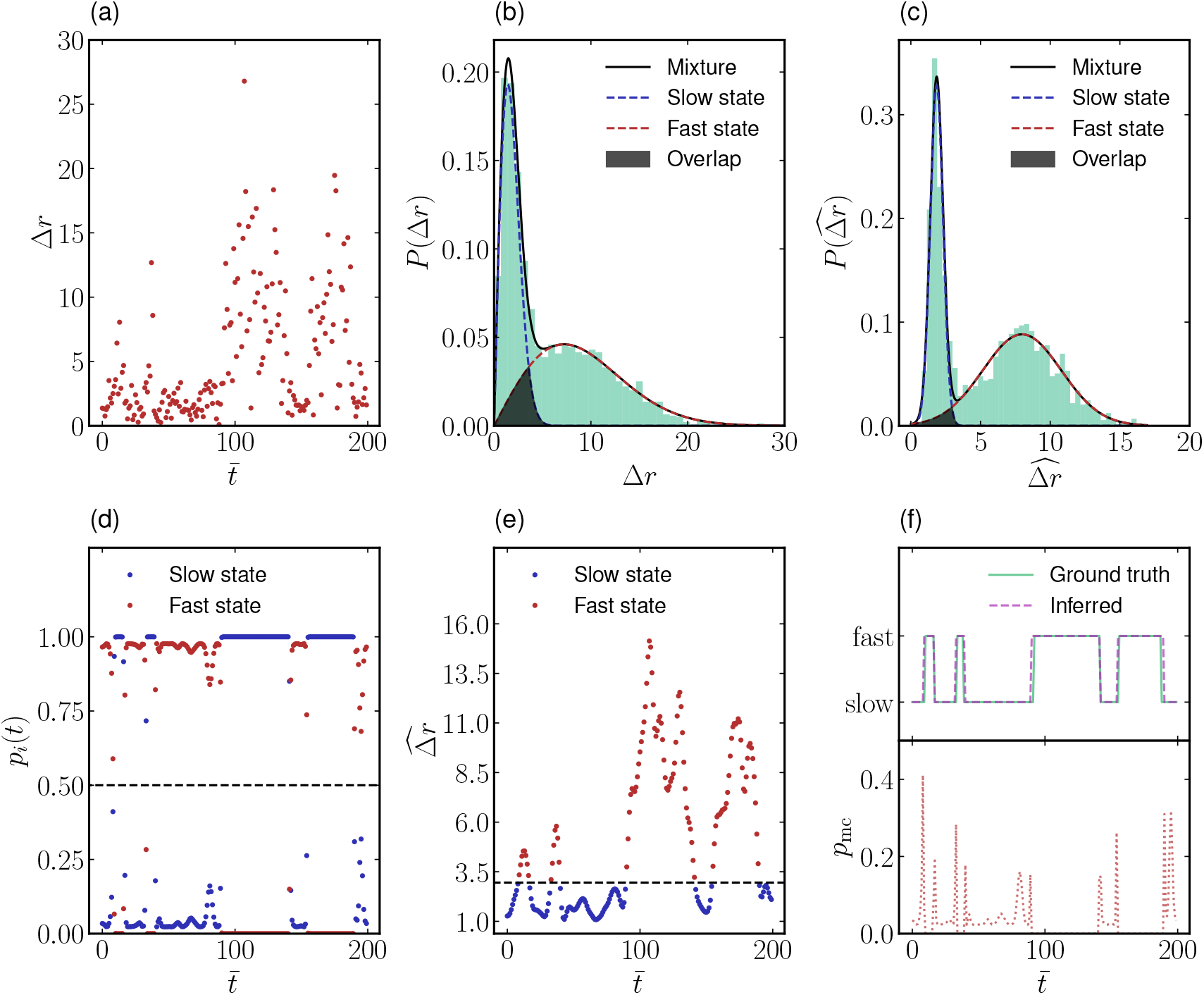
(a) Displacement time series of a 2-state Brownian particle, with 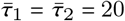, and *D*_2_/*D*_1_ = 5. (b) Displacement distribution of the *N*_*t*_ unfiltered trajectories fitted to a Rayleigh mixture according to Eq. (6). The overlap between the displacement distributions of individual states is shaded. (c) Displacement distribution of the same *N*_*t*_ trajectories, but now after filtering using the optimal filter width *f*\* fitted to a sum of two Gaussians, according to Eq. (8). (d) Inferred time series of the probability *p*_*i*_ of belonging to the diffusive state *i*, extracted from the filtered displacement distribution. The black dashed line indicates the threshold, corresponding to *p*_*i*_ = 0.5. (e) The displacement time series in (a) after filtering using the optimal filter width *f*\*. The dashed line corresponds to the optimised threshold between states 1 and 2, inferred by the intersection point between the two fitted Gaussians in (c). (f) Comparison between the ground truth and inferred state timeseries of the trajectory (top panel) and the inferred probability of misclassification, *p*_mc_ ≡ min (*p*_1_, *p*_2_)(bottom panel, *c*.*f*. Fig. 3d).

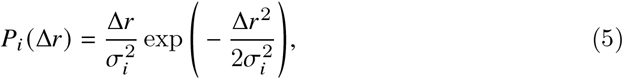

yielding a 2-state displacement distribution given by

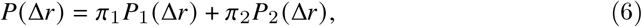

with the weights 𝜋_1_ = 𝜆_21_/(𝜆_12_ + 𝜆_21_) and 𝜋_2_ = 𝜆_12_/(𝜆_12_ + 𝜆_21_) such that 𝜋_1_ + 𝜋_2_ = 1, as shown in Fig. 3b.

From Figs. 3a and b, we observe that, although large displacements are more frequent in the state with the higher diffusion coefficient, small displacements are frequent in both states, leading to considerable overlap between the two probability densities. *D*_1_ and *D*_2_ can then be obtained by fitting the Rayleigh mixture model (RMM) in Eq. (6) to the measured distribution [16, 35]. However, the large overlap between the two probability densities makes a direct inference of the state at a given time point from the raw data challenging, and therefore an RMM cannot be used directly to detect transitions or infer values of 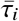. To reduce this overlap and enable an explicit segmentation of the trajectory into a time series of the state of a given particle, we convolute the displacement time series Δ*r* (*t*) with a discrete Gaussian filter 𝛽 of width *f* to obtain the *filtered* displacement time series 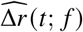:

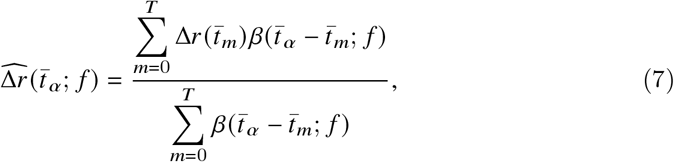

where 𝛽(*t*; *f*) = exp [−*t*^2^/(2 *f* ^2^)]. Here, we also tacitly assume that all state lifetimes 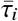 are longer than, or at least comparable to, the filter width *f* to avoid significant blurring of the state transitions.

This filtering procedure is a weighted, moving average, where the weights are given by a Gaussian distribution centered at each data point. In practice, this means that data points closer to the target point contribute more strongly to the filtered value, while points farther away contribute progressively less. For *f* ≪ Δ*t*, 𝛽 approaches a delta function, leading to a filtered displacement time series identical to the unfiltered one. Conversely, for *f* ≫ Δ*t*, 𝛽 yields an effectively constant displacement time series. For intermediate values, the temporal correlations introduced by the filtering process will reduce the standard deviation of *P*_*i*_ (Δ*r*), leading to a decrease in the overlap between the individual probability distributions *P*_1_ and *P*_2_. This suggests that the overlap 𝜃_12_ between *P*_1_ and *P*_2_, which we use as an effective cost function, is minimised for some optimal value *f*\* of the filter width. For values of *f* in the neighborhood of *f*\*, each point in the filtered displacement time series will therefore be influenced by only a small number of adjacent points in time, and the correlations introduced by the filtering are weak. In this case, the filtered sequence has enough independence that the central limit theorem applies, and 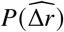 can be accurately described by a sum of two Gaussian distributions 𝒩 (𝜇, 𝜎), according to

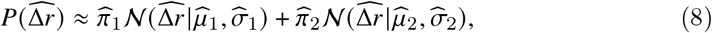

with 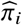 denoting the weights of the filtered displacement distribution for each state after the filtering process, and 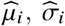,the corresponding means and standard deviations, where 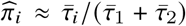 and we generally expect 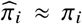 since the weights of the underlying two states are not altered by the filtering process. This decomposition allows us to use Gaussian mixture models (GMMs) for fitting the composite distribution. A GMM is a form of soft clustering that models data as a mixture of Gaussian distributions, which, unlike hard clustering methods that assign each data point to a single cluster without accounting for any uncertainty in cluster membership, compute the probability *p*_*i*_ that a given data point (or displacement) belongs to each component (state) *i* in the mixture. The parameters of the model — the means, covariances, and weights — are estimated using the expectation-maximization (EM) algorithm, which iteratively updates the probabilities of belonging to a given state and adjusts the model parameters to obtain an optimal fit [55, 56]; in our code, we use the GMM implementation provided by the `GaussianMixture`class in the `scikit-learn`library [57]. GMMs are employed to fit the distribution 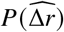 and to provide a pointwise estimate of the likelihood *p*_*i*_ that a given displacement originates from a specific diffusive state *i*. This fitting procedure further enables an approximation of the overlap 𝜃_12_ between the two distributions, expressed as

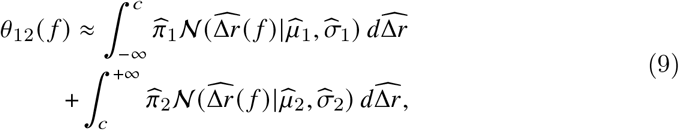

where 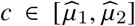 is the intersection point of the two weighted distributions, corresponding to the point where the probabilities of each of the two states are equal. For systems with multiple states, this generalises to the pairwise sum of the overlap, 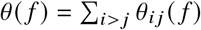. Fig. 3c shows the filtered version of the displacement distribution in Fig. 3b, which visually demonstrates the radically increased separability between the two states.

After finding *f*\*, we obtain the time series representing the relative probability of each data point belonging to either of the two diffusive states, as illustrated in Fig. 3d. By assigning each filtered displacement to the state associated with the highest probability, we retrieve a segmented time series of the filtered displacements (Fig. 3e). Additionally, the pointwise probability values serve as quantitative indicators of the confidence associated with the state classification at each time point. In Fig. 3f, we show the resulting, inferred time series of the two diffusive states, together with the “ground truth” transition points used when generating the trajectory. In spite of the relatively large overlap between the unfiltered distributions (Fig. 3b), the reconstructed time series in Fig. 3f shows that all transitions are correctly detected within a small time window, indicating a successful segmentation of the trajectory. Here, we have also show the time series of *p*_mc_ ≡ min(*p*_1_, *p*_2_) inferred from Fig. 3d, which measures the *a priori* probability of misclassification at a given time point. The possibility to temporally monitor this measure provides a fast estimate of the quality of the data and the filtering. In the following section, we will quantify the accuracy of this segmentation and its dependence on the parameters of the problem.

### Segmentation accuracy

In the following, we will demonstrate our method on 2-state Brownian trajectories generated as described in the previous section. Our main focus is on the discriminability between two states, quantified by the ratio of the two diffusion coefficients, 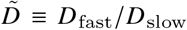 and the shortest of the two lifetimes, 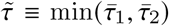, measured in units of the shutter time Δ*t*. The effect of the trajectory length, *T*, will also be examined, as well as the motion blur due to the limited shutter speed, as emulated by the averaging in Eq. (3), and the rescaled tracking error 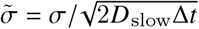.

We however start in the absence of blurring and tracking error, corresponding to 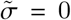 and *n* = 1 in Eq. (3), and investigate the accuracy of the segmentation for various values of 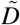 and 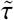. We define the accuracy as the proportion of time steps at which the inferred diffusive state matches the ground truth:

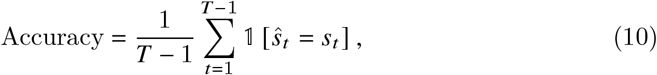

where Ŝ_*t*_ is the inferred diffusive state, *s*_*t*_ is the ground truth state at time *t*, and 𝟙 [·] is the indicator function, which equals 1 if the prediction is correct and 0 otherwise. Parameter space was explored by generating *N*_*t*_ = 100 trajectories of length *T* = 1000 for each pair (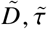). These parameters are important as 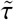 puts an upper bound on the range of possible filter size values before the filtering process starts to blur the two states, while 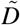 dictates the ideal filter size needed to minimise the overlap between the two states in the case of very large 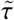.

Fig. 4a shows that our method gives a high precision segmentation (>90% accuracy) for a large range of (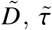), although becomes inaccurate for a small separation between the diffusion coefficients (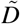 ≤ 3) or fast transition times (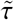 ≤ 20) compared to the camera resolution. A trade-off between 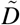 and 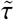 is observed, as small values of 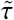 require a larger value of 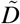 and vice versa in order to effectively minimise the cost function and obtain an accurate segmentation. Figures 4b and 4c highlight the underlying optimisation algorithm behind the segmentation method, respectively showing the overlap 𝜃_12_ and the accuracy functions of the filter width *f*. First of all, we note that 𝜃_12_ is a convex function across the whole domain of interest, allowing for a well-posed optimisation problem. Secondly, there is a strong anticorrelation between the overlap and the segmentation accuracy for all parameter values shown, which supports our use of overlap as a cost function. Although the maximum in accuracy is not perfectly aligned with the minimum in 𝜃_12_, their colocalisation is strong enough to ensure a near-optimal performance in spite of the simplicity and fully automated nature of the optimisation method. The results further show that the optimal filter width is generally small, falling in the range 1 ≤ *f* ≤ 5, meaning that it is generally well-separated from the smallest transition time 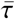 apart from the smallest values explored in Fig. 4a.

**Fig. 4.**
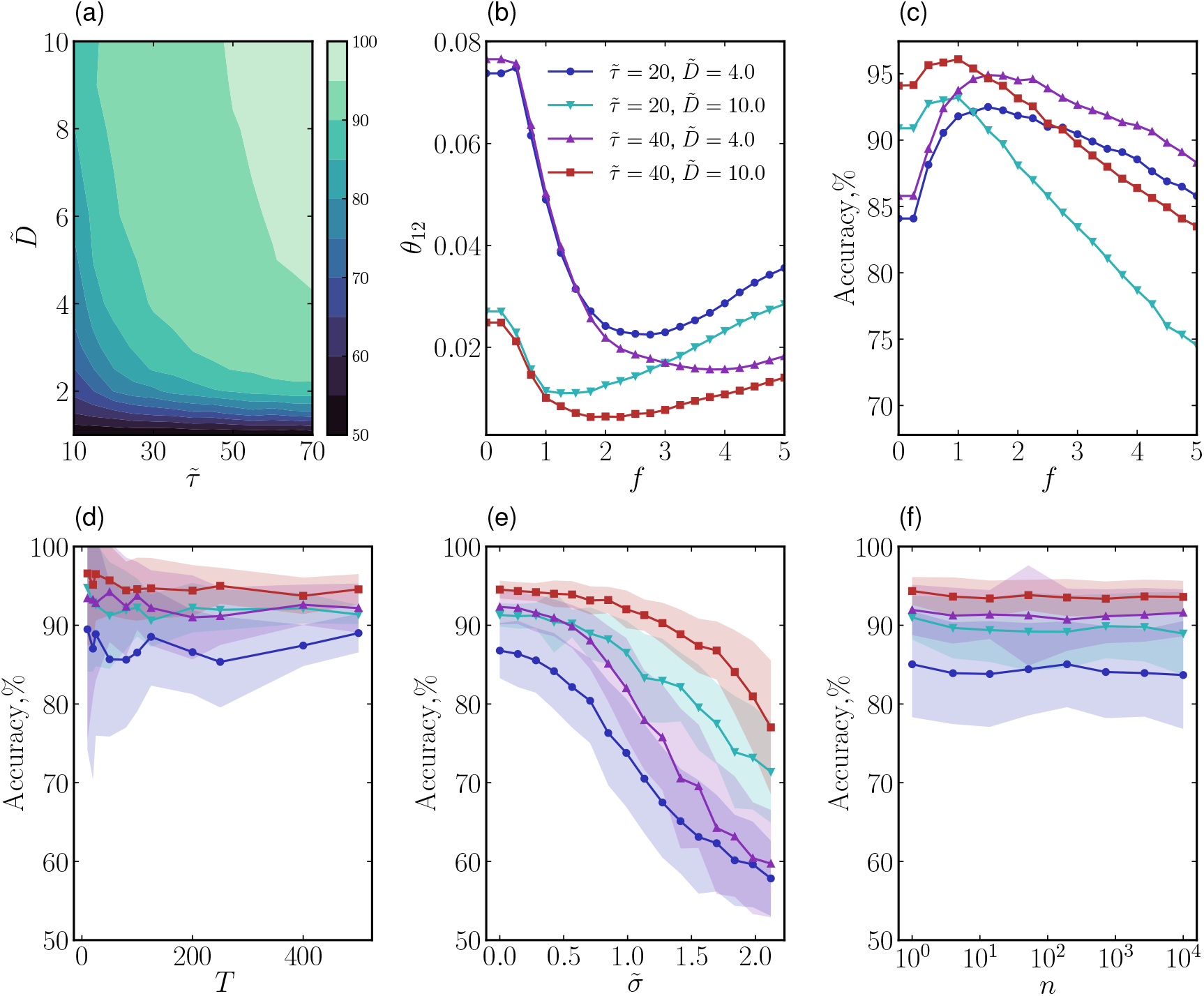
(a) Colour plot showing the segmentation accuracy as a function of 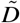 and 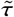, obtained after averaging over *N*_*t*_ = 100 trajectories of length *T* = 2000. (b) The overlap between the displacement distribution of the two states 𝜃_12_ and (c) the segmentation accuracy, both as a function of the filter width *f* for the same values of 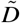 and 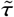 as in the other panels. (d) The accuracy as a function of the trajectory length *T* for various 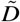 and 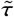, keeping fixed the total number of sampled displacements *N*_*t*_ × *T* = 20000. (e) The accuracy as a function of the localisation error 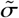 for the same values of 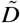 and 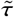 as in panel (b). (f) The accuracy as a function of the intensity of motion blur, as mimicked by varying *n* in Eq. (3). In this case, the “ground truth” when determining the segmentation accuracy for the blurred trajectory was determined through a majority vote of all *n* points involved in the blurring process. Where applicable, shadings represent one standard deviation of the accuracy over *N*_*t*_ = 100 trajectories.

In Fig. 4d we furthermore demonstrate that the method is independent of the length of individual trajectories *T*. To verify this, we generate *N*_*t*_ trajectories of varying lengths *T*, keeping the total number of displacements *N*_*t*_ × *T* constant, fixed at 20000 displacements. We observe that the accuracy of the segmentation method remains consistent regardless of trajectory length. While this might seem surprising, it is a direct consequence of the probabilistic nature of the segmentation method, which only relies on sampling a probability distribution of displacements between adjacent time points as in Figs. 3b and c, a process which does not require long individual time series. Therefore, the only strict requirement on the time series length for successful segmentation is that *T* ≫ *f*\*. However, as we will see in the following section, a reliable estimate of the lifetimes 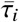 using the segmented trajectories impose more conservative conditions on the trajectory lengths.

We now add the effect of localisation error and motion blur by generating *N*_*t*_ = 200 Brownian trajectories with *T* = 1000, 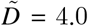 and 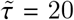, to which we apply different intensities of both spatial localisation error, as controlled by 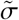, and motion blur, adjusted by changing *n*, the number of discretised steps over which the particle position in the original trajectory is averaged, when generating the trajectory according to Eq. (3). In Fig. 4e, we vary 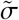 from 0, corresponding to no localisation error, to 2.1, corresponding to a case where the localisation error is significantly larger than the motion of the particle in its slowest state. The results show that the segmentation accuracy is essentially unaffected by the localisation error for 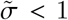. However, when 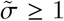, *i*.*e*., when the localisation error is of the order of the instantaneous displacement, the accuracy decays slowly, as expected. This is a consequence of the large noise introduced by the tracking error, which makes the overlap 𝜃_12_ larger and thus the states become harder to distinguish from each other.

In Fig. 4f, we show the corresponding impact of the motion blur as quantified by the number of points *n* used to generate the rescaled trajectory in Eq. (3). Qualitatively, this yields a spurious correlation between adjacent displacements 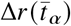 and 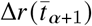 [54], which is generally challenging to handle using HMM-based methods [37]. For our segmentation algorithm, the main effect is a slight blurring of the unfiltered probability distribution in Fig. 3b, which however does not lead to any measurable decline in the accuracy of segmentation, as shown in Fig. 4f. Unlike the localisation error 𝜎, a direct translation of the parameter *n* to experimentally relevant ones is however not straightforward, as the underlying, discretised Brownian process used for the averaging is in itself not a faithful description of the real-world, continuous-time dynamics in the experimental system. Therefore, a continuous-time trajectory with motion blur corresponds to the limit *n* → ∞. For a full assessment of the impact of motion blur, a variation of the “blur factor”, *R*, which describes the impact of the shutter function on the trajectory is therefore more appropriate, although also more technically complicated. The data generation in Eq. (3) gives equal weight to each position in the underlying trajectory, corresponding to a top-hat (or square-wave) shutter function that counts every photon during the acquisition time, yielding *R*= 1/6 [54].

### Extraction of dynamical properties

Once the trajectories have been segmented, the diffusion coefficient for each state can be directly determined from the corresponding trajectory segments. Since the segmented trajectories correspond to *single-state* ones, the dynamical analysis is significantly simplified, allowing for simple extraction of model parameters such as diffusion coefficients and state lifetimes. Assuming isotropic, Brownian diffusion, the displacements in each dimension can be utilised to compute the diffusion coefficient by averaging over both dimensions according to [54]

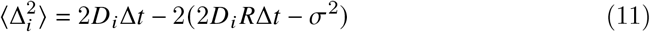

where 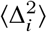 is the mean-squared, unidimensional displacement averaged over the displacements corresponding to the *i*^th^ state and the blur factor *R*equals 1/6 if all frames are weighted equally during the acquisition process [54], as in Eq. (3).

To verify the accuracy of the parameter estimation, in Fig. 5a and b we extract the diffusion coefficient from 200 segmented trajectories with *T* = 5000, 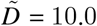, and 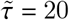 using Eq. (11). Fig. 5a shows that, even for low values of the localisation error, the faster of the two diffusion coefficients is systematically underestimated by ~ 10%. This is due to an asymmetry in the misclassification, where by a larger part of the classification mistakes misclassifies displacements from the “slow” state as being “fast” than vice versa, resulting in a slight understimation of *D*_fast_. On the other hand, the accuracy in determining *D*_*i*_ is suprisingly robust to the localisation error 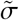 : even when this is in the region 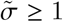where classification errors start becoming significant (green curve in Fig. 4e), the impact on the predicted diffusion is negligible until 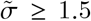, although its standard deviation (shaded regions in Fig. 5a) increases steeply with 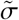, particularly for the slow state which gets the highest relative impact by the noise. This indicates that the additional classification errors introduced by localisation noise for moderate 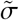 is symmetric in the sense that it leads to “fast” states being misclassified as “slow” ones as often as the opposite case, creating an effective cancellation of errors when calculating the diffusion coefficients. Fig. 5b, which shows the effect of motion blur on the extraction of *D*_*i*_, exhibits the same systematic underestimation of *D*_fast_, although the accuracy is largely unaffected by the introduction of motion blur even as *n* → ∞. Importantly, this does not imply that motion blur does not affect the displacement time series, but only illustrates that this effect can be efficiently corrected for using the additional term in Eq. (11), since the calculation of *D*_*i*_ is done on the single-state trajectories after segmentation.

**Fig. 5.**
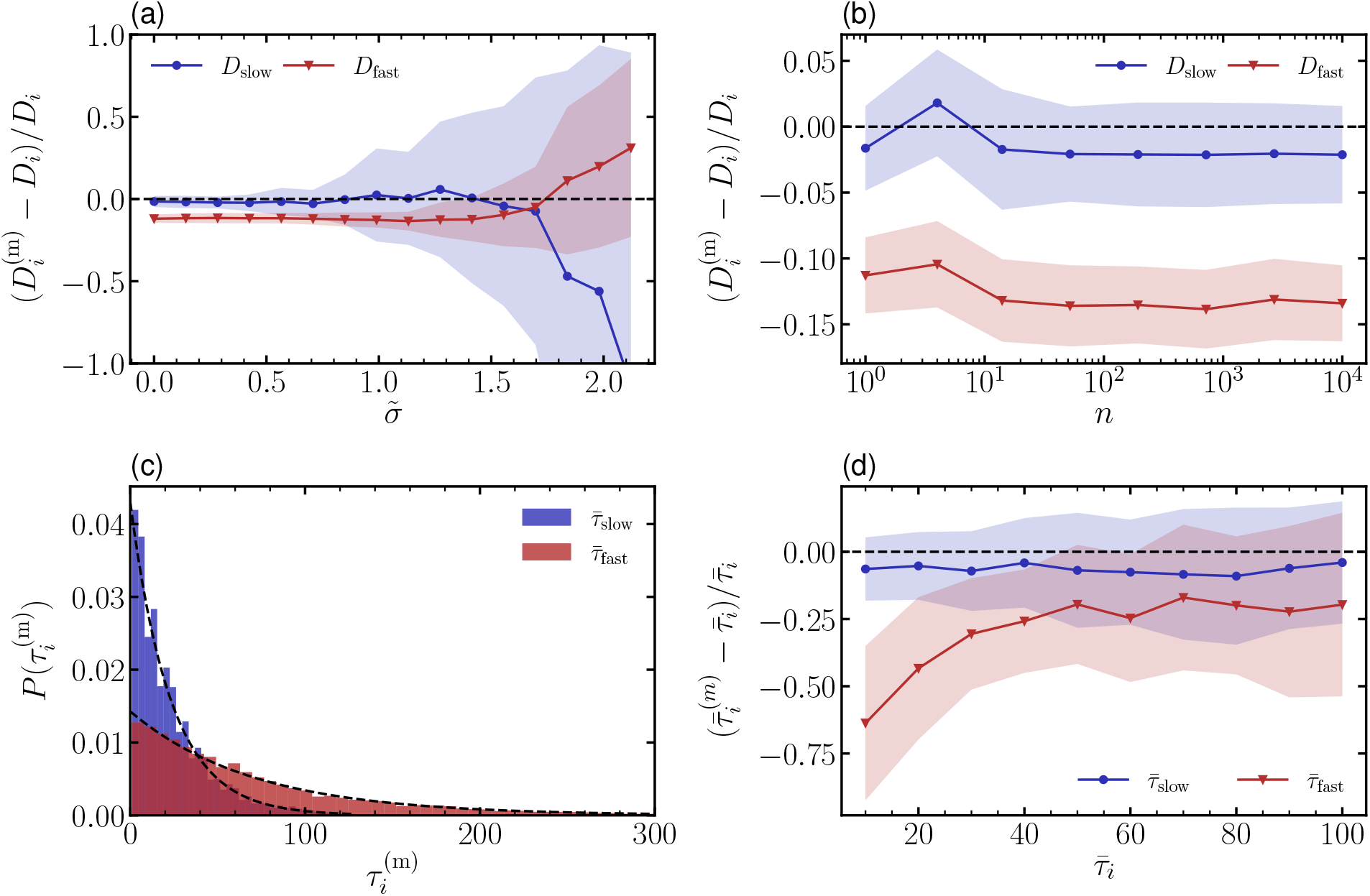
(ab) Mean relative error in the inferred diffusion coefficient 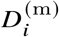, as a function of (a) the rescaled localisation error 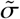 and (b) the blurring strength *n*, in both cases using 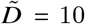 and 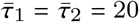. (c) Probability distribution of the measured lifetime 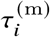. The distributions are fitted to an exponential (dashed lines), yielding estimated average lifetimes of 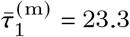 and 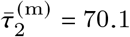, compared to the true values of 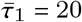 and 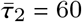, corresponding to a relative error of approximately 15%. (d) Relative mean error of the measured average lifetimes 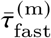 and 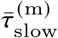 as a function of the true value 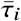, using 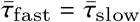. Where applicable, shading represents one standard deviation of the estimate measured over *N*_*t*_ = 200 trajectories.

Extracting the lifetimes of the two states is generally a much more demanding task, regardless of the segmentation method, as it intrinsically requires trajectories that are long enough to cover at least two transitions between the two states [50]. Similar to previous methods [58], we estimate the average lifetime 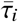 by assuming a Poissonian behaviour and sampling the lifetime 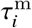 of segmented trajectories, followed by fitting of the resulting histogram to an exponential distribution 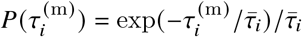, as shown in Fig. 5c. Compared to the determination of diffusion coefficients, this extraction is sensitive to errors in the segmentation process, since the misclassification of a displacement within a state introduces a “virtual transition”, leading to an underestimation of lifetimes, and primarily the lifetime 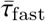 of the state with the faster diffusion. Nevertheless, Fig. 5d shows a relative error within ±25% for 𝜏 ≥ 30, consistent with the trend observed in Figure 4a, and indicating that the method can provide reasonably accurate lifetime estimates if using sufficiently numerous and long trajectories. This also illustrates another advantage compared to fitting the unfiltered distribution to a Rayleigh mixture model, which does not enable extraction of 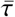, but only of the ratio between the two lifetimes.

### Experimental verification

The results in Figs. 3–5 all show examples of synthetic data, generated as discussed in the Model description section; in the following, we will illustrate the usability of our segmentation method also for experimental single-particle tracking data. To this end, we use a system consisting of Alexa Fluor 647-labeled, histidine-tagged SLAMF6 proteins (Catalog #9299-NT, R&D Systems) anchored to a supported lipid bilayer, following the protocol described by Dam *et al*. [59]; a representative microscopy image of this system is shown in Fig. 6a. Visual inspection of the motion of the labelled proteins (Fig. 6b) indicates the presence of two different protein ensembles, one exhibiting free, 2-dimensional diffusion in the membrane and one with highly restricted mobility. In spite of this, the unfiltered displacement distribution in Fig. 6c shows a single, broad peak with an extended tail. The distribution is furthermore relatively poorly fitted by two Rayleigh distributions; in spite of this, the filtered distribution (Fig. 6d) shows two well-separated, Gaussian peaks that are well-fitted by the GMM. The resulting diffusion coefficients, as calculated from Eq. (11) are *D*_1_ = 0.057 µm^2^ s^−1^ and *D*_2_ = 1.44 µm^2^ s^−1^, corresponding to 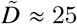. This large ratio between the two particle mobilities, together with the fact that the two populations remain essentially constant over the course of the experiment, makes this a relatively “easy” system to segment, and further verification on experimental systems with dynamic binding-unbinding kinetics is clearly warranted. Nevertheless, the significant difference between the filtered and unfiltered distributions in Fig. 6 clearly illustrates the radical effect of our, relatively simple, segmentation algorithm when applied to experimental data.

**Fig. 6.**
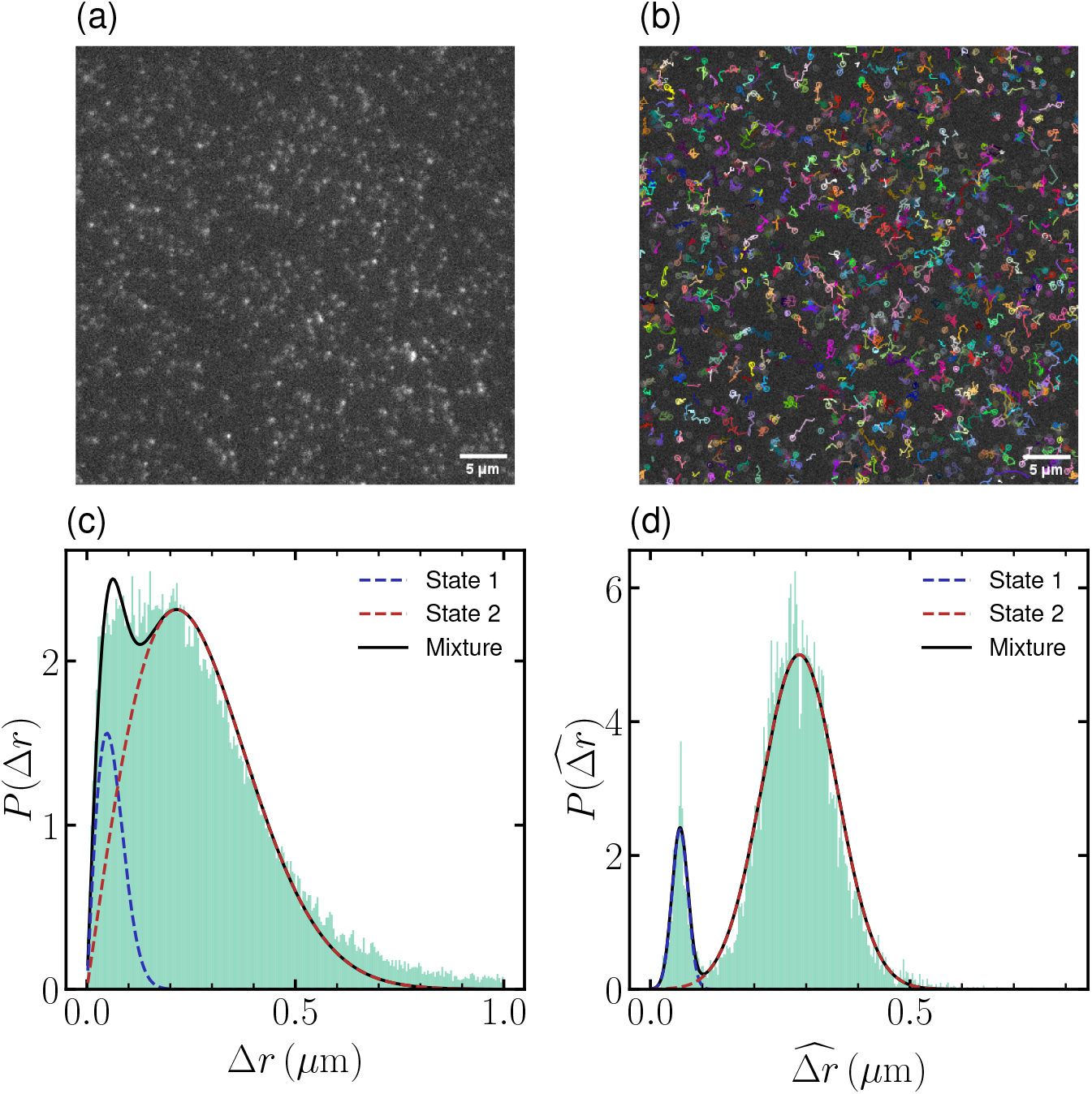
(ab) Single-molecule images of Alexa Fluor 647 labeled histidine-tagged SLAMF6 (Catalog #9299-NT, R&D Systems) anchored to a supported lipid bilayer, following the protocol described previously [59] (a) and with overlaid, coloured particle tracks (b). Images were acquired using total internal reflection fluorescence (TIRF) microscopy with a frame interval of 30 ms [59]. Particle detection and tracking were performed over 100 frames using TrackMate 8 [60] in Fiji [61]; tracks shorter than five frames were excluded and the maximum displacement between consecutive snapshots was set to 1.0 µm. For visualization purposes, tracks in panel (b) are limited to maximum 20 frames. (c) Unfiltered displacement distribution calculated from the experimental trajectories shown in (b), with a fitted sum (solid line) of two Rayleigh distributions (dashed lines) corresponding to *D*_1_ = 0.058 µm^2^ s^−1^ and *D*_2_ = 1.16 µm^2^ s^−1^ after correcting for motion blur. Note the relatively poor quality of the fit both in the tail and at the peak. (d) Filtered displacement distribution obtained by applying a Gaussian filter with an optimal filter size chosen to minimize overlap between the two diffusive states, yielding diffusion coefficients *D*_1_ = 0.057 µm^2^ s^−1^ and *D*_2_ = 1.44 µm^2^ s^−1^. Note that the filtering procedure renders two distinct, Gaussian peaks with much better fits than for the unfiltered data in panel (c).

## Discussion

In this study, we have presented and verified a novel method for the segmentation of multi-state Brownian trajectories using an automated optimisation of a Gaussian filtering process. By minimising the overlap between the displacement distributions of different diffusive states, our method provides a robust and interpretable framework for trajectory segmentation in cases where the two states are (*i*) sufficiently separable in terms of diffusion coefficients (*D*_fast_/*D*_slow_ ≥ 4) and (*ii*) sufficiently long-lived compared to the camera resolution (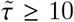). Our results show that the method performs with high accuracy under experimentally realistic conditions, where it can be 10 used to quantitatively characterise the diffusive dynamics, and hence to infer, *e*.*g*., the kinetics of protein-ligand binding and the relative size of bound and unbound ligand populations, both in live cells [16] and in biophysical model systems [25]. Importantly, the method is furthermore both easy to implement and computationally efficient: the segmentation of *N*_*t*_ = 1000 trajectories each containing 100 steps requires only approximately 15 seconds on a standard laptop. The main strength of the proposed method therefore lies in it being accurate but simultaneously lightweight and transparent: the model does not require training, the trajectory segmentation is fast enough to be done on-the-fly, and the quality of the filtered probability distributions and corresponding fits can be directly evaluated visually or by goodness-of-fit estimates to assess the quality of the segmentation. It therefore also presents itself as a potentially attractive alternative for on-line preprocessing of experimental data, both in order to efficiently optimise experimental parameters, and to characterise the different diffusive populations (*e*.*g*., the ratio between mobile and immobile molecules), as illustrated in Fig. 6. Furthermore, the fact that the segmentation step does not require any information about the dynamical parameters such as diffusion coefficients or lifetimes, these can be inferred from the segmented, single-state trajectories, avoiding the complication of parameter fitting to multi-state trajectories.

Even though we verify our method on the specific case of Brownian trajectories with Markovian switching dynamics, this choice was made out of simplicity rather than to specifically suit the segmentation method. On the contrary, one strength of the method is that it makes no explicit assumption about the underlying nature of the switching dynamics or diffusive processes, since it only relies on the particles’ displacement between two adjacent points in time. After filtering, this displacement time series will result in Gaussian probability distributions as long as the filter is narrow enough that the correlations introduced by the filtering are weak, as illustrated in Fig. 6d. Thus, the filtering process is in no way specific to Brownian diffusion, in contrast, for example, to the fitting of Rayleigh mixtures to the (unfiltered) displacement PDF, as shown in Fig. 6c, which assumes an underlying Brownian process. However, for strongly non-Brownian statistics, the ensuing calculation of *D*_*i*_ using Eq. (11) would need to be generalised accordingly. Furthermore, the form of the switching dynamics plays no role in the segmentation process as long as the dwell time is significantly larger than the time resolution of the microscope, again since the segmentation algorithm only handles displacements between adjacent time points. However, the underlying switching dynamics will, by construction, affect the distribution of residence times extracted from the segmented data: For data with sufficiently long and numerous tracks, the dwell time probabilities shown in Fig. 5c can in principle be extracted from the segmented trajectories and analysed without any *a priori* assumption of their functional form. In spite of this flexibility, as we discuss in the following, there are several possible extensions of the method that could be implemented to handle the complexities present in experimental data from biological systems.

First of all, unlike some more advanced methods that can infer an optimal number of states *K*, our approach uses *K* as a fixed parameter which is known *a priori*. Inferring *K* can in principle be done using standard likelihood-based model selection [35, 38]. One possible such route would be to fit GMMs with *K* = 1, …, *K*_max_ states, and compute the Bayesian information criterion, BIC(*K*) = −2 log(*L*) +*K* log(*N*_*t*_ ×*T*), where *L* denotes the maximized likelihood under the *K*-state model [35]. The optimal number of states is then selected as the one that provides some physically motivated trade-off between goodness-of-fit and model complexity. While this algorithm is straightforward to implement, this procedure tends to overestimate *K* since the penalty of adding more states is much smaller than the decrease in the log-likelihood and therefore requires a somewhat *ad hoc* trade-off in the last step. A generalisation of our method to a *fixed K* >2 is however very straightforward, and a preliminary implementation indicates similar accuracies for 3-state trajectories as long as the diffusivities are well separated; hence, keeping *K* as a fixed (but user-defined) parameter probably provides the best trade-off between flexibility and automation.

Secondly, our method tacitly assumes *well-defined* diffusion constants *D*_*i*_ of each state, which includes systems with two well-separated ensembles or spatial regions with distinctly different dynamics. Although we used fixed values of *D*_*i*_ in our method verification, we expect the method to also work well for non-constant diffusivities with sufficiently narrow distributions so that the overlap between the two peaks remains small, although ML-or HMM-based methods [62] are likely more well-suited for systems with broad and overlapping distributions of *D*_*i*_.

In spite of these built-in limitations, we believe that the presented method provides a versatile and light-weight option for the analysis of complex displacement statistics from experimental particle trajectories. It therefore contributes to the growing portfolio of single-particle tracking methods by providing a computationally efficient and interpretable alternative to existing segmentation techniques for multi-state Brownian trajectories, forming another methodological tool in the toolbox for understanding the dynamics, molecular transport processes, and interactions in complex cellular systems.

## Data availability

The experimental data used to generate Fig. 6 can be downloaded from https://github.com/joakimstenhammar/GMM-Seg.

## Code availability

The software used for the generation of Brownian trajectories, segmentation analysis and calculation of dynamical properties is available for download at https://github.com/joakimstenhammar/GMM-Seg.

## Acknowledgments

Helpful discussions with Carlo Manzo and Giovanni Volpe are kindly acknowledged. This work was funded by the Knut and Alice Wallenberg Foundation (Grant 2019.0079). JS and TA acknowledge funding from the Swedish research council (Grants 2019-03718 and 2022-03475, respectively).

## Author Contributions

IEK, JML, TD, EC, TA, PJ, and JS conceptualised the research. IEK, JML, and JS designed the model. IEK and JML conducted the software implementation and data analysis. IEK, JML, and JS wrote the paper with input from all the authors. All authors have read and approved the final version of the manuscript.

